# Synthetic guide sequence to generate CRISPR-Cas9 entry strains in *C. elegans*

**DOI:** 10.1101/2025.06.16.659939

**Authors:** Karen I. Lange

## Abstract

CRISPR/Cas9 genome editing has become an important and routine method in *C. elegans* research to generate new mutants and endogenously tag genes. One complication of CRISPR experiments is that the efficiency of single-guide RNA sequences can vary dramatically. One solution to this problem is to create an intermediate entry strain using the efficient and well-characterised *dpy-10* guide RNA sequence. This “d10 entry strain” can then be used to generate your knock-in of interest. However, the *dpy-10* sequence is not always suitable when creating an entry strain. For example, if your gene of interest is closely linked to *dpy-10* on LGII or if you want to use the *dpy-10* as a co-CRISPR marker for the creation of the entry strain then you can not use the *dpy-10* sequence. This publication reports a synthetic guide sequence, GCTATCAACTATCCATATCG, that is not present in the *C. elegans* genome and can be used to create entry strains. This guide sequence is demonstrated to be relatively robust with a knock-in efficiency that varies from 1-11%. While this is lower than the efficiency observed with *d10* entry strains, it is still sufficient for most applications. This guide sequence can be added to the *C. elegans* CRISPR toolkit and is particularly useful for generating entry strains where the standard *dpy-10* guide sequence is not suitable.

## Description

Over the last decade, CRISPR/Cas9 genome editing has become an important and routine method in *C. elegans* research (Kim et al., 2022). There are many applications for genome editing including engineering specific mutations and endogenously tagging genes. In *C. elegans*, a co-CRISPR strategy using dominant phenotypic markers (Arribere et al., 2014) has been a very successful method to facilitate reproducible and reliable genome editing. The most widely used co-CRISPR gene is *dpy-10;* a specific *dpy-10* missense mutation, Arg92Cys, exhibits a dominant roller (Rol) phenotype while imprecise edits cause a recessive dumpy (Dpy) phenotype. Cas9 cutting of the *dpy-10* guide sequence is very efficient with homozygous Dpy or DpyRol worms frequently observed in the F1 generation of CRISPR edited worms (Arribere et al., 2014). One difficulty in designing new CRISPR experiments is that there is no method to accurately predict the efficiency of a guide RNA *a priori*. It has previously been described that you can take advantage of the efficient *dpy-10* guide sequence to create an entry strain for your gene of interest to facilitate genome editing with a single efficient guide RNA (El Mouridi et al., 2017). You can create a “d10 entry” strain by inserting *dpy-10* guide sequence at your region of interest and then use this strain in a second round of CRISPR genome editing to reliably introduce your desired edits (El Mouridi et al., 2017). There are many advantages to this d10 entry strain approach including increasing the reliability of low efficiency knock-in edits, easily editing the same gene multiple times, and the ability to engineer scarless edits. This method has proven to be effective as evidenced by the many publications that cite this strategy (Dietz et al., 2021; Lynch et al., 2022; Placentino et al., 2021; Schreier et al., 2022; Silva-García et al., 2019; Vigne et al., 2021). There are several situations where a d10 entry strain is not possible such as if your gene of interest is closely linked to *dpy-10* on chromosome II or if you want to use the efficient *dpy-10* co-CRISPR marker when you generate the entry strain. This publication reports an alternative guide sequence that can be used to generate reliable entry strains.

I wanted to knock-in mNeonGreen(mNG) at the N-terminus of *cep-290* and the closest adjacent PAM was 25 nucleotides from the start codon. After screening 4520 co-CRISPR positive F1 and not isolating the desired knock-in, I decided to attempt the “d10 entry” strain approach (El Mouridi et al., 2017). I designed an ssODN repair template that included a 25bp deletion and insertion of the d10 guide sequence on the antisense strand directly after the start codon (Figure 1A). The repair template also included a mutation in a nearby EcoRI cut site to facilitate detection of the edit. I screened 124 *unc-58* co-CRISPR positive F1 for the loss of the EcoRI site and identified one potential edit. This allele w as isolated named *cep-290(oq120)*. It was sequenced revealing that *cep-290(oq120)* differed significantly from the original repair template; the d10 sequence was partially inserted and there was no adjacent PAM (Figure 1A). I observed that a novel PAM site was unexpectedly generated in the *cep-290(oq120)* allele so it could potentially function as an entry strain. This new PAM has a guide sequence, GCTATCAACTATCCATATCG, that is a hybrid of the *cep-290* and the *dpy-10* sequences so I refer to it as a “synthetic guide”. I reasoned that this sequence may have high on-target efficiency because it exhibits some of the qualities that have been previously reported in efficient Cas9 guide sequences including a high GC content (40%), multiple CA or AC dinucleotides, and a G in the 20 position (Doench et al., 2014; Wong et al., 2015). To check for off target sites in the *C. elegans* genome, I employed the CRISPR-Cas9 guide RNA design checker that is available on the Integrated DNA Technologies website. All identified off targets have at least 4 mismatches (Figure 1B). By targeting the synthetic guide sequence in the *cep-290(oq120)* entry strain I was able to efficiently generate the mNG::*cep-290* strain by screening only 184 co-CRISPR positive F1s and isolating two knock-in lines.

**Figure 1.**
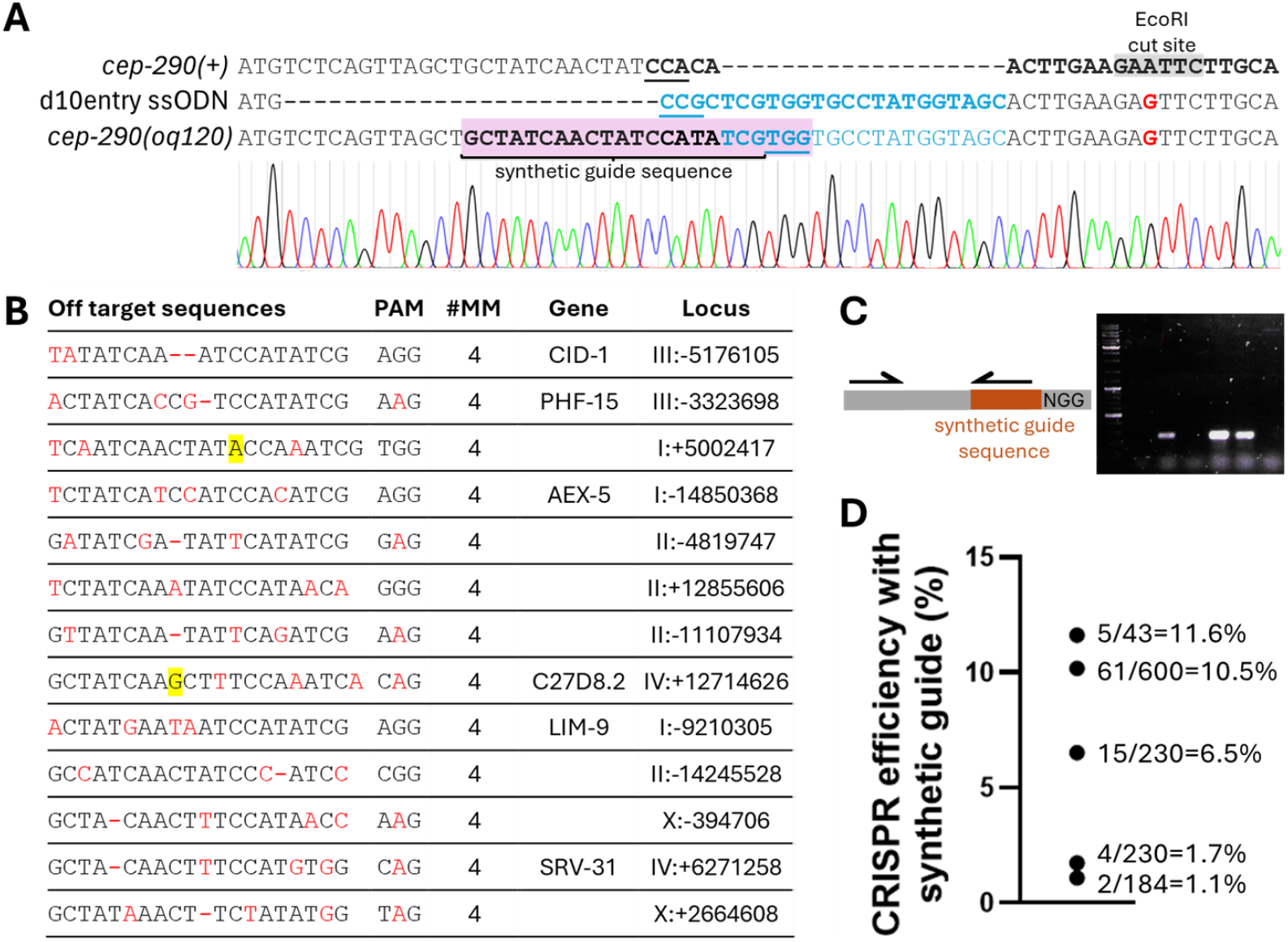
Synthetic guide sequence generated when attempting to generate a d10 entry strain at the 5’ end of *cep-290*. **A)** DNA sequences of wild-type *cep-290*, the repair template for the designed “d10 entry” allele, and the actual allele isolated. The repair template (ssODN) was designed to introduce a 25 bp deletion and insertion of the d10 guide sequence directly after the start codon. The *cep-290(oq120)* allele (chromatogram shown) was detected and isolated due to the missense mutation that disrupts an EroRI site (red), however the sequence was not consistent with the engineered entry strain. A new guide with PAM (highlighted in purple) was generated in this allele. This sequence has partial homology with *cep-290* and *dpy-10* and we have termed it a synthetic guide sequence. Relevant guide sequences are bolded and PAMs are underlined. **B)** Predicted off target effects of the synthetic guide sequence identified with the CRISPR-Cas9 guide RNA design checker available on the Integrated DNA Technologies website. All identified off targets have at least 4 mismatches. Mismatches are red. Insertions are highlighted in yellow. **C)** Insertion of the synthetic guide sequence can be detected efficiently by a 2 primer PCR reaction where one primer is gene specific and the other binds to the synthetic guide sequence. A product will only be amplified if the synthetic guide sequence has been inserted in the genome. A sample gel is shown. **D)** Knock-in efficiency using this synthetic guide sequence at various loci in the *C. elegans* genome ranges from approximately 1-11%. Efficiency was calculated by taking the number of PCR positive F1 pools and dividing it by the total number of F1 that were screened.

I have since used this synthetic guide sequence to make multiple entry strains and found it to be reliable and efficient. Insertion of the entry strain can be easily detected in F1 progeny using a 2 primer PCR reaction with one gene specific primer and one primer that is complementary to the synthetic guide sequence (Figure 1C). The efficiency of knock-in strains generated with this guide has ranged from 1% to 10% (Figure 1D). This is lower than the reported efficiency of d10 entry strains which ranged from 3-19% (El Mouridi et al., 2017) but is still sufficient for use in generating entry strains. Interestingly, both of these guide sequences exhibited a wide range of efficiencies when inserted at different genomic loci; this observation highlights how CRISPR efficiency can be affected by non-sequence specific factors such as chromatin state (Horlbeck et al., 2016; Isaac et al., 2016).

The synthetic guide sequence reported here can be used in any *C. elegans* CRISPR application where you might use an entry strain. I have found it to be particularly useful in situations where the d10 guide sequence is not suitable, such as if the gene of interest is linked to *dpy-10* on chromosome II or when you want to use *dpy-10* as the co-CRISPR marker while generating the entry strain. The synthetic guide sequence described here was generated by accident and not designed. It may be possible to rationally design a more efficient synthetic guide sequence, but since the sequence described here is functional it may not be worth the time and resources that would be required to attempt to improve it. In conclusion, this sequence is another useful option that can be added to the *C. elegans* CRISPR tool kit.

## Methods

### Nematode strains

*Caenorhabditis elegans* strains were maintained at 20°C on NGM agar plates seeded with *E. coli* (OP50) using standard worm maintenance techniques (Brenner, 1974; Stiernagle, 2006).

The following worm strains were used or generated in this study: N2 wild-type, OEB931 *cep-290(oq120[entry strain]) I*, and OEB932 *cep-290(oq121[mNeonGreen::cep-290]) I*.

### PCR to generate *mNG::cep-290* repair template

mNeonGreen(mNG) is licensed by Allele Biotechnology and Pharmaceuticals (Shaner et al., 2013). *C. elegans* codon optimised mNeonGreen (Hostettler et al., 2017) was amplified from a plasmid, dg353 (a gift from D. Glauser), and a 12 amino acid flexible linker (GTGGGGSGGGGS) was added to the 3’ end of the mNG sequence as previously described (Lange et al., 2021). 35 base pair homology arms were added to the mNG sequence with two rounds of high-fidelity PCR as per manufacturer’s instructions (Velocity, BIO-21098, Meridian BioScience).

### Generation of entry strains *and mNG::cep-290* with CRISPR

CRISPR experiments were performed by microinjection of the Cas9 ribonucleoprotein complex (Paix et al., 2015). CRISPR reagents were purchased from IDT: Alt-R Cas9 Nuclease V3 (IDT, #1081058), Alt-R tracrRNA (IDT, #1072533), and custom synthesised gene specific Alt-R crRNA. All RNA for CRISPR experiments was reconstituted with 5 mM Tris (pH 7.5) and stored at −75°C. Single-stranded oligonucleotides (ssODN) repair templates were ordered from Sigma-Merck and reconstituted with 1 M Tris pH 7.4 and kept at −20°C. CRISPR mixes were prepared as previously described (Lange et al., 2021) and incubated at 37°C for 15 min prior to microinjection into the gonads of young adult hermaphrodites. A co-CRISPR approach with *dpy-10* or *unc-58* was used (Arribere et al., 2014); F1 progeny with the co-CRISPR marker phenotype were pooled in groups of 3-8 worms and edits were detected by PCR. The *d10::cep-290* entry allele was identified by loss of an EcoRI cut site; restriction digests were performed with EcoRI-HF (R3101S, NEB) as per the manufacturer’s instructions. Subsequent entry strains were identified by using a primer that was complementary to the synthetic guide sequence. Sanger sequencing by Eurofins Genomics was used to determine the sequence of all CRISPR alleles generated.

### Calculating efficiency of the synthetic guide RNA sequence

Knock-in efficiency was calculated by dividing the number of PCR positive F1 pools by the total number of co-CRISPR positive F1 that were screened.

## Reagents

### crRNA*

*cep-290:* TGCAAGAATTCTTCAAGTTG

Synthetic guide: GCTATCAACTATCCATATCG

*dpy-10:* GCTACCATAGGCACCACGAG

*unc-58:* ATCCACGCACATGGTCACTA

*For convenience crRNA sequences are shown as their corresponding DNA sequences.

### ssODN repair templates

*d10::cep-290* ssODN: GCACCTTTTTACTAGCACAAATGTACTGAGACATGCCGCTCGTGGTGCCTATGGTAGCACTTGAAGAGTTCTTGCAAAATGATGGTCCTACCGAGGAAGAAGT

*dpy-10* ssODN: CACTTGAACTTCAATACGGCAAGATGAGAATGACTGGAAACCGTACCGCATGCGGTGCCTATGGTAGCGGAGCTTCACATGGCTTCAGACCAACAGCCTAT

*unc-58* ssODN: GTGGTATAAAATAGCCGAGTTAGGAAACAAATTTTTCTTTCAGGTTTTTCTGTCGTTACCAT GTGCGTGGATCTTGCGTCCACACATCTCAAGGCGTACTT

### Primers to generate *mNG::cep-290* repair template

For: GCACCTTTTTACTAGCACAAATGTACTGAGACATGGTGTCGAAGGGAGAAGAGG Rev: TCTTCCTCGGTAGGACCATCATTTTGCAAGAACTCTTCAAGTTGTGGATAGTTGATAGCAGCTAACTGAGATCCGCCACCTCCAG

Nested For: GCACCTTTTTACTAGCACAAATG Nested Rev: TCTTCCTCGGTAGGACCATC

### Genotyping and sequencing primers for *cep-290(oq121)*

mNG Rev: AGGCTCCATCCTCGAATTGC For: CTGTCAGTTTCTCATGGTGC

Rev: ATCCTCTGCCTCCTTGGAC Seq: TCCACCCTCCTACACACTC

### Synthetic guide specific primer

Rev: CGATATGGATAGTTGATAGC

## Acknowledgements

I would like to acknowledge Oliver Blacque for supporting this project which began when was a postdoctoral researcher in his group. The *C. elegans* optimised mNeonGreen plasmid was a gift from Dominique Glauser (University of Fribourg, Switzerland). Some worms were provided by the Caenorhabditis Genetics Center, which is funded by NIH Office of Research Infrastructure Programs (P40 OD010440).

## Author Contributions

KIL was responsible for conceptualization, investigation, validation, visualisation, writing, and editing.

